# Transcription Factor EB regulates phosphatidylinositol-3-phosphate levels on endomembranes and alters lysosome positioning in the bladder cancer model

**DOI:** 10.1101/2020.07.10.196931

**Authors:** Pallavi Mathur, Camilla De Barros Santos, Hugo Lachuer, Bruno Latgé, François Radvanyi, Bruno Goud, Kristine Schauer

## Abstract

Lysosomes orchestrate degradation and recycling of exogenous and endogenous material, thus controlling cellular homeostasis. Little is known how this organelle changes during malignant transformation. We investigate the intracellular landscape of lysosomes in a cellular model of bladder cancer. Employing standardized cell culture on micropatterns we identify a phenotype of peripheral lysosome positioning prevailing in bladder cancer but not normal urothelium. We show that lysosome positioning is controlled by transcription factor EB (TFEB) and that lysosomal dispersion results from TFEB activation downstream of lysosomal Ca^2+^ release. Remarkably, we find that TFEB regulates phosphatidylinositol-3-phosphate (PtdIns3P) levels on endomembranes which recruit FYVE-domain containing proteins, such as the motor adaptor protrudin, for anterograde movement of lysosomes. Altogether, we uncover lysosome positioning as result of PtdIns3P activation downstream of TFEB as a potential biomarker for bladder cancer. Moreover, we reveal a novel role of TFEB in regulating cellular PtdIns levels, conceptually clarifying the dual role of TFEB as regulator of endosomal maturation and autophagy, two distinct processes controlled by PtdIns3P.

**Statement of significance:** Here we provide the first atlas for the landscape of the lysosomal compartment in bladder cancer and reveal the mechanistic role of TFEB in regulating endosomal PtdIns3P levels and subsequent lysosomal dispersion. We unveiled lysosomal positioning as a potential biomarker for malignant bladder cancer which might arise as an actionable target for cancer therapy.

## Introduction

Accelerated cellular division and enhanced motility are pathological characteristics of malignant cells both leading to an increase in energetic demand. More than being the ‘stomach’ of eukaryotic cells for nutrient acquisition, late endosomes/lysosomes (referred to as lysosomes hereafter) have emerged as a cellular hub for metabolism and signaling (Ballabio and Bonifacino, 2020; Lawrence and Zoncu, 2019; Perera and Zoncu, 2016; Perera et al., 2019; Thelen and Zoncu, 2017) and play an important role during cancer development (Hämälistö and Jäättelä, 2016; Perera and Zoncu, 2016). Lysosomes are morphologically heterogeneous acidic compartments that are functionally similar to yeast and plant vacuoles. They are specialized in the degradation of extracellular molecules and pathogens internalized by endocytosis or phagocytosis, as well as the intracellular recycling of macromolecules and organelles sequestered by autophagy. In addition to the orchestration of cellular clearance, lysosomes play an important role in cellular nutrient availability controlled by the serine/threonine kinase complex of mammalian target of rapamycin complex 1 (mTORC1) (Saxton and Sabatini, 2017). Active mTORC1 assembles at the surface of lysosomes through the integration of chemically diverse nutrient and growth factor signaling to promote protein biosynthesis (Ballabio and Bonifacino, 2020; Saxton and Sabatini, 2017; Thelen and Zoncu, 2017). Conversely, absence of nutrients triggers the dissociation and inactivation of mTORC1 and consequently to the activation of downstream catabolic pathways. Active mTORC1 targets MiT/TFE transcription factors, including transcription factor EB (TFEB) and MITF, that are both master regulators of lysosome biogenesis and autophagy (Settembre et al., 2011). MiT/TFE transcription factors have been implicated in the development of cancer, including renal cell carcinoma, pancreatic adenocarcinoma, sarcoma and melanoma, MITF being an important oncogene in melanoma (Perera et al., 2019). Moreover, it has been shown that TFEB overexpression as well as a positive feedback mechanism between mTORC1 and TFEB was sufficient to promote cancer growth in mouse models (Calcagnì et al., 2016; Di Malta et al., 2017).

Although lysosomes are important for nutrient acquisition and the regulation of metabolism, both prerequisites for malignant growth, little is known how lysosomes change during cancer development. Here, we compare the intracellular landscape of the lysosomal compartment in a collection of bladder cancer cell lines to normal human urothelium (NHU). Bladder cancer represents one of the most frequently-diagnosed cancer types worldwide and is among the most common neoplasms in men in North America and Europe, thus representing an important health burden(Antoni et al., 2017). Bladder carcinomas are highly diverse and are classified into non-muscle-invasive bladder cancers (NMIBC) and muscle-invasive bladder cancers (MIBC) with luminal-like and basal-like subtypes (Choi et al., 2014; Rebouissou et al., 2014). NMIBC are often papillary (stage Ta) and low-grade but show a high recurrence rate (60%). Alternatively, NMIBC can be carcinoma in situ (CIS, stage Tis) showing flat lesions and frequent progression to invasive cancers (T1). MIBC are classified by stages T2-T4 and high-grade malignant transformation. Investigating the normal and pathologic landscape of lysosome positioning in cells representing different stages of bladder cancer, we here reveal organelle-level deregulation in malignant cells and identify TFEB as major regulator of phosphatidylinositol-3-phosphate (PtdIns3P) homeostasis in this context.

## Results

### High-grade bladder cancers are characterized by the peripheral positioning of lysosomes

Because of the importance of lysosomes in cellular homeostasis and their role in promoting cancer progression, we aimed at a systematic analysis of lysosome morphology in a panel of genetically diverse bladder cell lines in comparison to primary normal human urothelium (NHU) cells. We have analyzed the broadly studied bladder cancer cell lines RT4, MGHU3, RT112, KU19-19, T24, TCCSup and JMSU1 that represent the diversity of bladder carcinomas (Zuiverloon et al., 2018). RT4, MGHU3, RT112 represent low-grade, luminal cancers of the papillary subtype, whereas KU19-19 represents high-grade, basal cancers and T24, TCCSup and JMSU1 represent high-grade cancers of mixed subtypes (Warrick et al., 2016; Zuiverloon et al., 2018). To compare these different cells at the morphological level, we cultured them on identical crossbow-shaped micropattern substrates. All tested cells were fully spread after 3 h of incubation, visualized by the average projection of the actin cytoskeleton (**Fig. S1A**), indicating that all cells adapted well to the micropatterns. We visualized the lysosomal compartment in all cells by immunofluorescence staining of the lysosomal-associated membrane protein 1 (LAMP1/CD107a) (**Fig. 1A**). Images were acquired in 3D and lysosomes were segmented to obtain quantitative information of their spatial organization, volume and numbers per cell. To visualize the average lysosome organization, we plotted 3D density maps (Duong et al., 2012; Schauer et al., 2010b) representing the smallest cellular volume that contains 50% of lysosomes (**Fig. 1B**). Notably, while in NHU cells lysosomes were positioned centrally, they were found to be spread out to the periphery in cancerous cells with the strongest phenotype exhibited in high-grade lines (**Fig. 1A, B**). Because the total cell area is standardized by the micropattern and thus identical in all cells, we calculated the nearest neighbor distance (NND) of lysosomes in each cell. Concomitantly, whereas the average NND in low-grade RT4 and MGHU3 cells was not significantly different from NHU cells, those of all other analyzed cell lines was significantly increased (**Fig. 1C**), indicating that lysosomes are more scattered in these cells. No clear trend in the number of lysosomes per cell (**Fig. 1D**) or average volume (**Fig. 1E**) was found among the tested cell lines. However, lysosomal volume negatively correlated with lysosomal number (**Fig. S1B**), indicating that few large lysosomes are in balance with many small ones. Principal component analysis (PCA) of the transcriptome data of these cells indicated that replicates of the NHU clustered together and separately, and that RT4 and MGHU3 were the most different compared to the other cell lines (**Fig. S1C**). Comparison between selected low-grade MGHU3 (luminal-type and central lysosomes) and RT112 (luminal-type and scattered lysosomes), and high-grade KU19-19 (basal-type) and JMSU1 cells (mixed-type) in invasion assays into collagen matrix from spheroids revealed, as expected, that MHGU3 was the less invasive cell line (invasion at 5 d), followed by RT112 (invasion at 3 d), KU19-19 and finally JMSU1 that both invaded at 1 d with different efficiency (**Fig. S1D**). To verify that changes in lysosomal positioning were not induced by micropatterning, we additionally analyzed lysosomes in classical cell culture conditions in selected cell lines. We measured the averaged squared distance of lysosomes to the center of mass of the cell (statistical inertia) normalized to the cell size (**Fig. S1E,F**). In agreement with our density map and NND analysis, the lysosome dispersion significantly increased from MGHU3 to JMSU1 cells. Our analyses collectively indicate that the lysosomal compartment shows differences between NHU and bladder cancer cell lines. Whereas some low-grade bladder cancer cell lines reveal central lysosomes similar to NHU cells, high-grade bladder cancer cells are characterized by a scattered, peripheral positioning of the lysosomal compartment.

**Figure 1.**
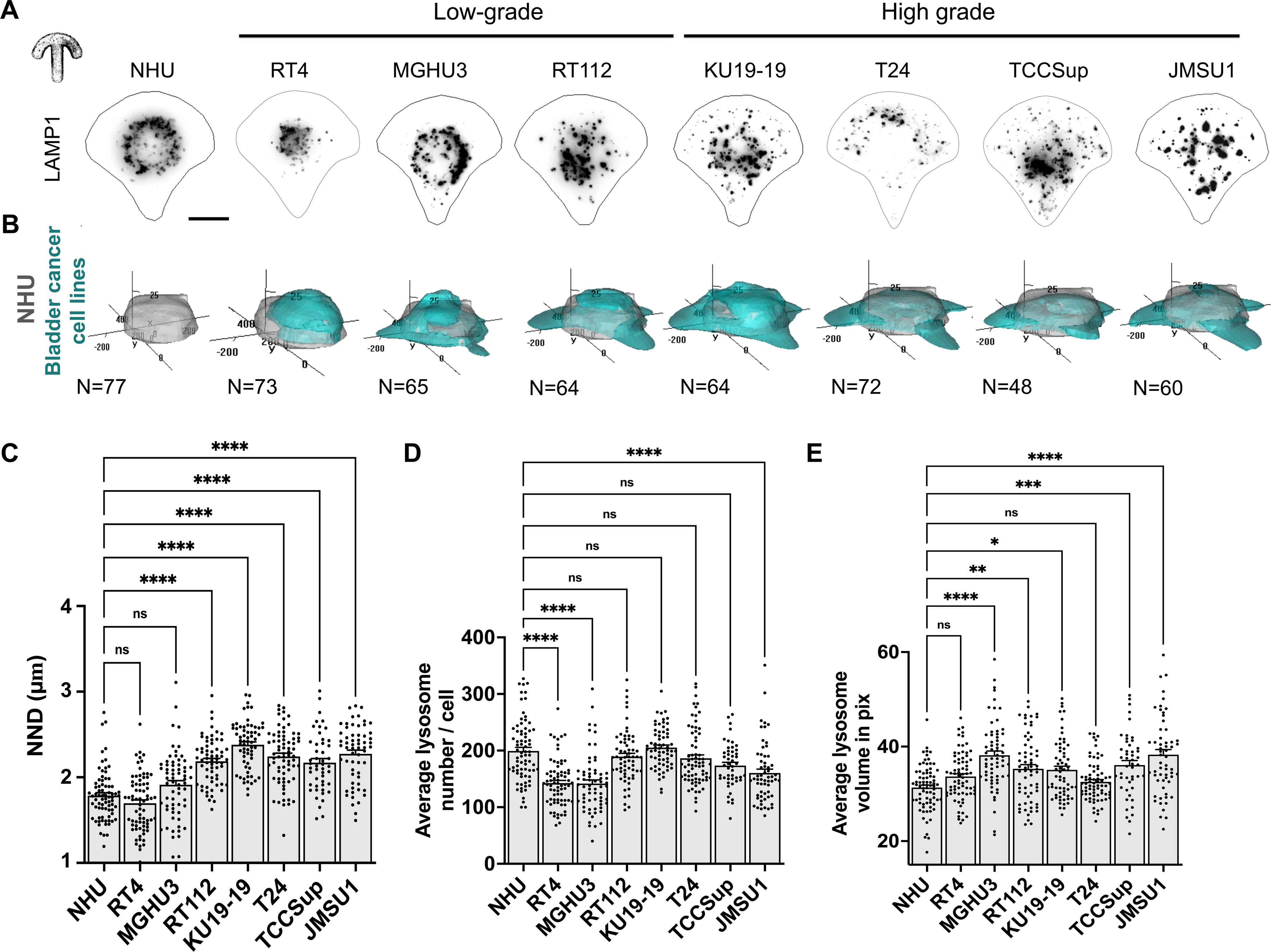
High-grade cancer cell lines are specifically characterized by scattered, peripheral positioning of lysosomes. **A**. Representative images of lysosomes visualized by immunofluorescence staining against the <u>l</u>ysosomal-<u>a</u>ssociated <u>m</u>embrane <u>p</u>rotein 1 (LAMP1/CD107a) in normal human urothelium (NHU) and bladder cancer cell lines RT4 (ATCC^®^ HTB-2™), MGHU3 (Lin et al., 1985), RT112 (Marshall et al., 1977), KU19-19 (Tachibana et al., 1995), T24, TCCSup (Nayak et al., 1977), JMSU1 (Morita et al., 1995) cells cultured on crossbow-shaped adhesive micropatterns for better comparison. Scale bar is 10 μm. **B**. 3D probabilistic density maps of lysosomes of n cells of NHU, MGHU3, RT112, KU19-19 and JMSU1. The 50% contour visualizes the smallest cellular volume containing 50% of lysosomes. **C**. Nearest neighbor distance (NND) between lysosomes in NHU (n=76), RT4 (n=73), MGHU3 (n=65), RT112 (n=64), KU19-19 (n=64), T24 (n=72), TCCSup (n=48) and JMSU1 (n=60). Adjusted p-values of testing against NHU condition are RT4: >0.9999, MGHU3: 0.1943; RT112: <0.0001; KU19-19: <0.0001; T24: <0.0001; TCCsup: <0.0001; JMSU1: <0.0001 in a Kruskal-Wallis test with Dunn’s test for multiple comparisons; ns p >0.1 and **** p < 0.0001, error bars are SEM. **D**. Average numbers of lysosomes per cell in NHU (n=76), RT4 (n=73), MGHU3 (n=65), RT112 (n=64), KU19-19 (n=64), T24 (n=72), TCCSup (n=48) and JMSU1 (n=60). Adjusted p-values of testing against NHU condition are RT4: <0.0001; MGHU3: <0.0001; RT112: >0.9999; KU19-19: 0.8807; T24: >0.9999; TCCsup: 0.2068; JMSU1: <0.0001 in a Kruskal-Wallis test with Dunn’s test for correction for multiple comparisons; ns p > 0.1 and **** p < 0.0001, error bars are SEM. **E**. Average volume of lysosomes in NHU (n=76), RT4 (n=73), MGHU3 (n=65), RT112 (n=64), KU19-19 (n=64), T24 (n=72), TCCSup (n=48) and JMSU1 (n=60). Adjusted p-values of testing against NHU condition are RT4: 0.1414; MGHU3: <0.0001; RT112: 0.0048; KU19-19: 0.0110; T24: >0.9999; TCCsup: 0.0003; JMSU1: <0.0001 in a Kruskal-Wallis test with Dunn’s test for multiple comparisons; ns p > 0.1, * p < 0.1, *** p < 0.001 and **** p < 0.0001, error bars are SEM.

### Dispersed lysosomes reveal alterations in the mTORC1-TFEB signaling axis

Lysosomes are the cellular signaling platform for the mammalian target of rapamycin complex1 (mTORC1), a main regulator of metabolisms, proliferation and survival. Because mTORC1 is regulated by lysosomes positioning (Korolchuk et al., 2011; Perera and Zoncu, 2016), we tested whether altered lysosome landscape across different bladder cancer cell lines affected mTORC1 signaling. First, we analyzed mTORC1 localization by co-visualizing mTOR and LAMP1 by immunofluorescence and measuring the fraction of mTOR that localized on the LAMP1-positive compartment. We found that about 15-20 % of mTOR signal was found on lysosomes. Although RT112 showed slightly but significantly more mTOR on lysosomes, the levels of mTOR on lysosomes were comparable between the tested cell lines (**Fig. 2A, B**). Next, we tested mTORC1 activity monitoring the phosphorylation of the direct downstream substrate p70-S6 Kinase 1 (S6K1). Surprisingly, we found that less S6K1 was phosphorylated in high-grade as compared to low-grade cells although total S6K1 levels were similar in all cell lines (**Fig. 2C and S2A**). As expected, the mTORC1 inhibitor rapamycin (Dumont and Su, 1995; Liu et al., 2010) as well as starvation decreased S6K1 phosphorylation in all cell lines confirming mTORC1 specificity (**Fig. S2B**). Next, we tested another important mTORC1 substrate, the transcription factor EB (TFEB), which appears as a novel player in carcinogenesis (Calcagnì et al., 2016; Di Malta et al., 2017). We transfected cells with TFEB-EGFP and monitored its localization in cells 72 h post transfection. Whereas TFEB-EGFP showed cytosolic localization in MGHU3 and RT112 cells (40% of mean TFEB-EGFP intensity was found in nucleus), more than 70% of the mean intensity of TFEB-EGFP was found in the nucleus of KU19-19 and JMSU1 cells (**Fig. 2D, E**). This nuclear localization indicated hyperactivation of TFEB in high-grade bladder cancer cells. Indeed, inspection of the gene expression of known TFEB-regulated genes such as *RAGD* and *TSC1* revealed an increase in the expression of these genes in high-grade as compared to low-grade cells (**Fig. S2C**). Inhibition of mTORC1 by rapamycin in low-grade RT112 led to a translocation of TFEB-EGFP to the nucleus (**Fig. 2F,G**) consistent with mTORC1-specific TFEB-EGFP phosphorylation. Contrary, activation of mTORC1 by U-18666 to stimulate phosphorylation of TFEB (Davis et al., 2021) in high-grade JMSU1 cells did not change TFEB-EGFP localization (**Fig. S2D, E**). It has been shown that the nuclear translocation of TFEB is additionally regulated through dephosphorylation by calcineurin (Medina et al., 2015). Calcineurin is activated by cytosolic calcium that is released from the lysosomes via mucolipin-1, also known as transient receptor potential cation channel, mucolipin subfamily, member 1 (TRPML1). Thus, we inhibited mucolipin-1 in high-grade bladder cancer cells using GW-405833 (ML-SI1) or incubated cells with the calcium chelator BAPTA for 2 h each. Both treatments led to the cytoplasmic translocation of TFEB-EGFP (**Fig. 2H, I**) indicating that increased dephosphorylation of TFEB by calcineurin in response to lysosomal calcium release strongly contributes to the nuclear accumulation of TFEB in high-grade bladder cancer cells. Together, our results indicate that peripheral lysosome positioning in high-grade KU19-19 and JMSU1 cells correlates with differences in the mTORC1-TFEB nutrient signaling pathway as compared to low-grade cell MGHU3 and RT112 although similar levels of mTORC1 recruitment on lysosomes was observed between the different grade cell lines. Moreover, our results depict an increased nuclear localization of TFEB in high-grade KU19-19 and JMSU1 cells, thus, a possible transactivation of TFEB regulated genes.

**Figure 2.**
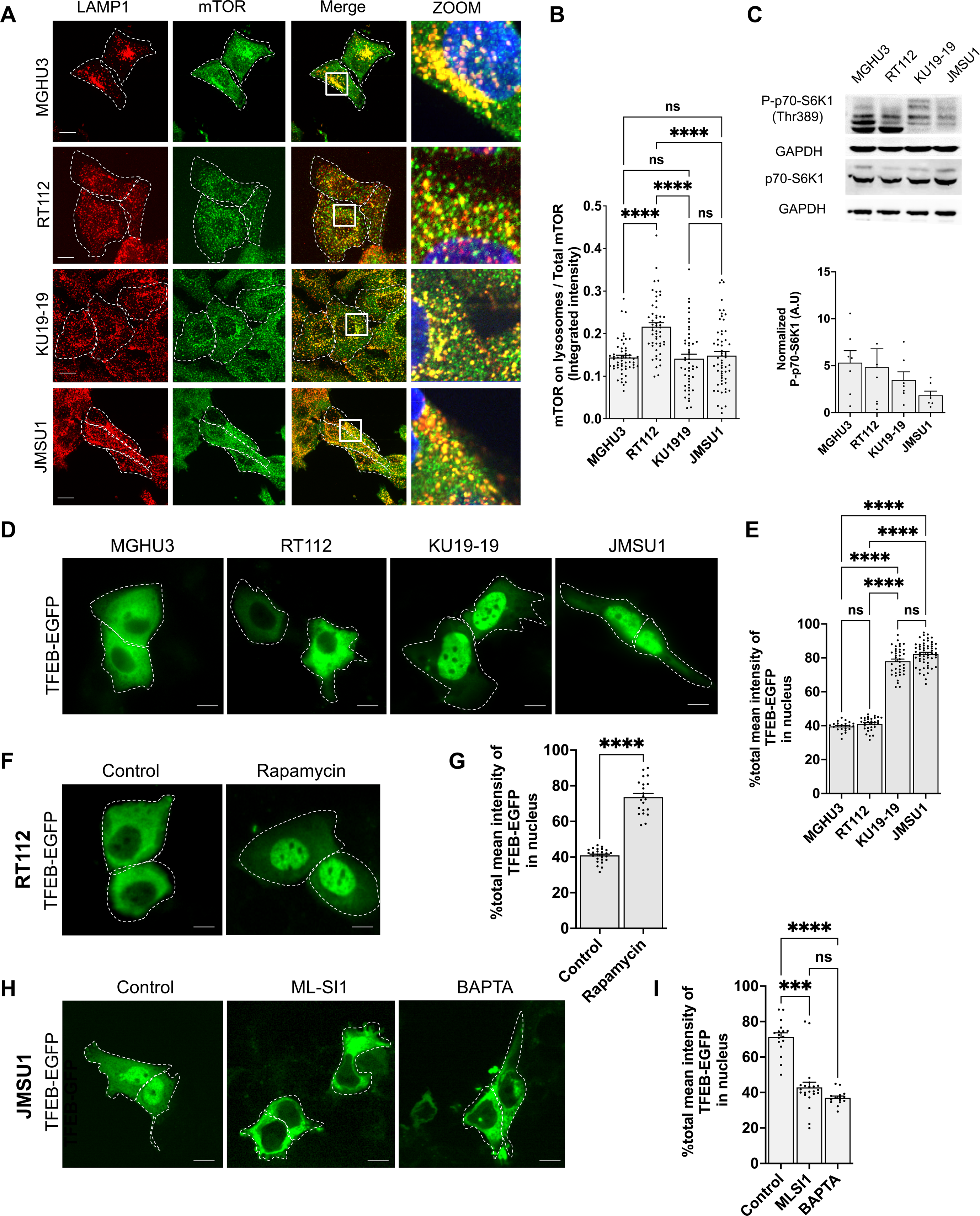
Dispersed lysosomes reveal alterations in the mTORC1-TFEB signaling axis. **A**. Immunofluorescence staining of the lysosomal-associated membrane protein 1 (LAMP1, CD107a) and mTOR in MGHU3, RT112, KU19-19 and JMSU1. The zoom shows the merged image for both proteins in the white box. Scale bars equal 15 μm **B**. Quantification of mTOR intensity on lysosomes normalized to total cellular mTOR (approximately 50 cells for each cell line; **** p <0.0001; Kruskal-Wallis test with Dunn’s test for multiple comparison, error bars are SEM. **C**. Western blot analysis of phosphorylated p70-S6 Kinase 1 (P-p70-S6K1 Thr389) and total p70-S6K1 as well as GAPDH loading control in MGHU3, RT112, KU19-19 and JMSU1 cells and quantification of P-p70-S6K1 levels from n=7 experiments, error bars are SEM. **D**. Representative images of MGHU3, RT112, KU19-19 and JMSU1 cells transfected with TFEB-EGFP for 72 h. Scale bars equal 10 μm. **E**. Quantification of the nuclear fraction of the total mean TFEB-EGFP fluorescent intensity in MGHU3 (n=23), RT112 (n=31), KU19-19 (n-39) and JMSU1 cells (n=57). **** p <0.0001; Kruskal-Wallis test with Dunn’s test for multiple comparison. Data are depicted as mean ± SD. **F**. Representative images of RT112 cells transfected with TFEB-EGFP for 72 h and treated with 10 μM rapamycin for 2 h. Scale bars equal 10 μm. **G**. Quantification of the nuclear fraction of the total mean TFEB-EGFP fluorescent intensity in control and rapamycin conditions (for n>20 cells in each condition). **** p <0.0001; Mann-Whitney test. Data are depicted as mean ± SD. **H**. Representative images of JMSU1 cells transfected with TFEB-EGFP for 72 h and treated with ML-SI1 or BAPTA AM for 3 h. Scale bars equal 10 μm. **I**. Quantification of the nuclear fraction of the total mean TFEB-EGFP fluorescent intensity in control, ML-SI1 and BAPTA AM treatment conditions (for >15 cells in each condition). *** p<0.001 and **** p <0.0001; Mann-Whitney test. Data are depicted as mean ± SD.

### Lysosome positioning changes are under the control of TFEB in bladder cancer cells

It has been previously reported that TFEB regulates lysosomal positioning (Willett et al., 2017), thus, we investigated whether increased nuclear translocation of TFEB in bladder cancer cell lines could lead to peripheral lysosome positioning. First, we tested whether stimulating nuclear translocation of TFEB in low-grade bladder cancer cells (RT112) triggered peripheral lysosome positioning. Cells were treated with rapamycin to induce TFEB nuclear translocation (**Fig. 2F,G and 3A**) and lysosomes were visualized by immunofluorescence against LAMP1. Inspection of classically cultured cells revealed recurrent accumulation of lysosomes at the cell periphery (**Fig. 3A**). To quantify this, we cultured cells on adhesive micropatterns and calculated the nearest neighbor distance (NND) of lysosomes in RT112 cells. Lysosomes were more dispersed after rapamycin treatment in micropatterned cells (**Fig. 3B**) and the average NND was significantly increased as compared to untreated controls (**Fig. 3C**). To specifically test the role of TFEB, we next targeted this member of the MiT/TFE family via RNA interference in high-grade (JMSU1) cells where TFEB is mostly nuclear. Silencing of TFEB by either a pool of four siRNAs or four individual siRNAs significantly reduced TFEB protein levels after 3 d and reversed the scattered lysosome phenotype in high-grade JMSU1 cells (**Fig. 3D and S3A-C**). Quantification on micropatterns revealed a significant decrease in the average NND of lysosomes (**Fig. 3E, F**) confirming TFEB-dependent regulation of lysosomes in these high-grade bladder cancer cells.

**Figure 3.**
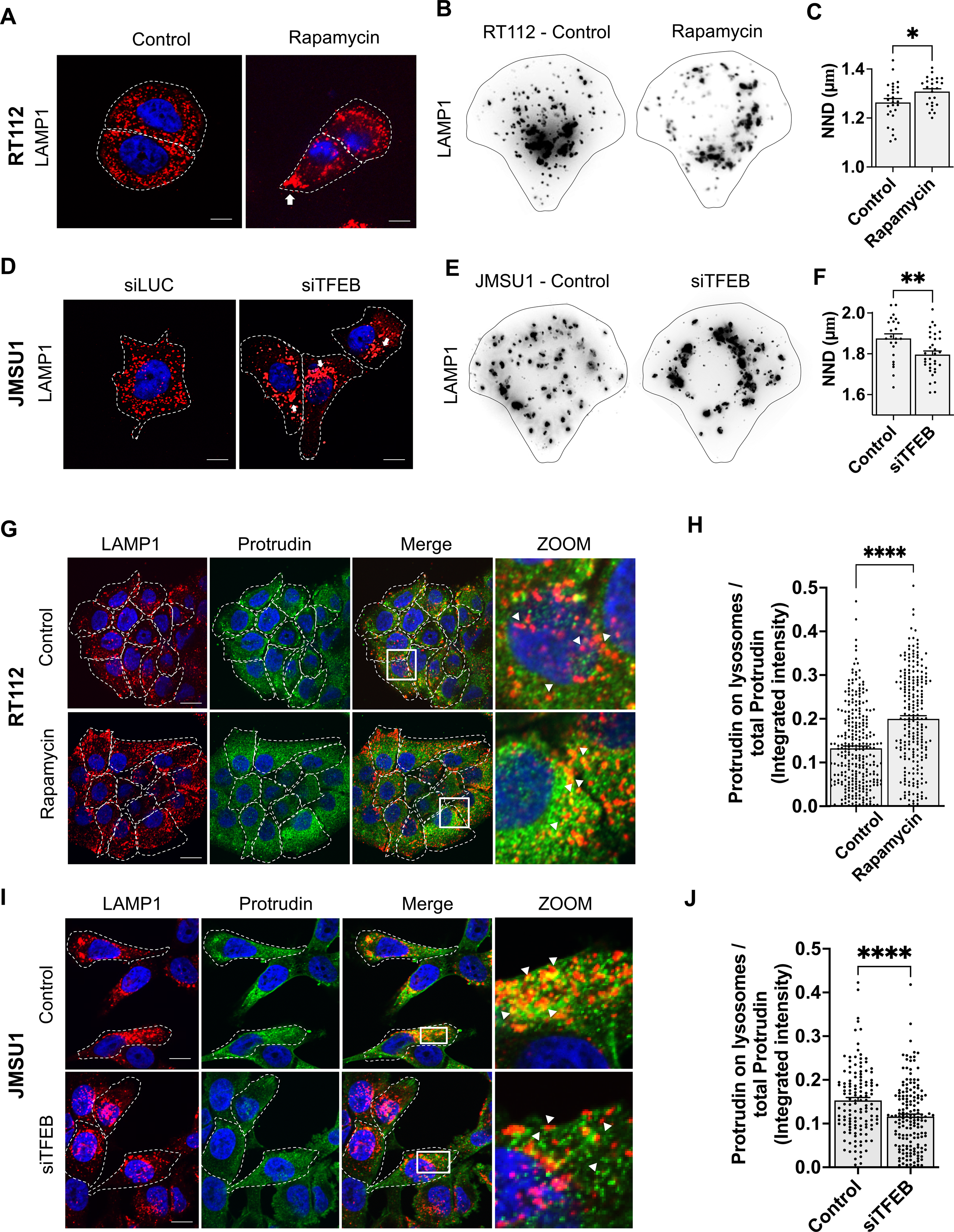
Lysosome positioning changes are under the control of TFEB in bladder cancer cells. **A**. Immunofluorescence staining of the lysosomal-associated membrane protein 1 (LAMP1, CD107a) in control (DMSO) and rapamycin (10 μM) treated RT112 cells. White arrow shows the peripheral clustering of lysosomes. Scale bars equal 10 μm. **B**. Representative images of lysosomes visualized by immunofluorescence staining against LAMP1 in micropatterned RT112 cells in control and rapamycin treatment. **C**. Nearest neighbor distance (NND in μm) between lysosomes in micropatterned control (n=25) and rapamycin treated (n=27) RT112 cells; * p <0.05 in a Mann-Whitney U test, error bars are SEM. **D**. Immunofluorescence staining of LAMP1 in JMSU1 cells treated with siLUC and siTFEB for 72 h. White arrow shows the perinuclear clustering of lysosomes. Scale bars equal 10 μm. **E**. Representative images of lysosomes visualized by immunofluorescence staining against LAMP1 in micropatterned JMSU1 cells in control and siTFEB treatment conditions. **F**. Nearest neighbor distance (NND in μm) between lysosomes in micropatterned control (n= 23) and siTFEB (n= 34) treated JMSU1 cells; ** p < 0.005 in a Mann-Whitney U test, error bars are SEM. **G**. Immunofluorescence staining of LAMP1 (red) and protrudin (green) in control (DMSO) and rapamycin (10 μM) treated RT112 cells. Zoom shows the merged image of both proteins in the white box. White arrow shows the colocalization between LAMP1 and protrudin. Scale bars are 15 μm. **H**. Quantification of protrudin integrated intensity on lysosomes normalized to total cellular protrudin, in 290 control and 227 rapamycin treated RT112 cells; ****p<0.0001 in a Mann-Whitney U test, error bars are SEM. **I**. Immunofluorescence staining of LAMP1 (red) in and protrudin (green) in JMSU1 cells in control (siLUC) and siTFEB (72 h) treatment conditions. Zoom shows the merged image of the two proteins in the white box. White arrow shows the colocalization between LAMP1 and protrudin. Scale bars are 15 μm. **J**. Quantification of protrudin integrated intensity on lysosomes normalized to total cellular protrudin, in 131 control (siLUC) and 167 siTFEB JMSU1 cells; **** p<0.0001 in a Mann-Whitney U test, error bars are SEM.

It has been shown that lysosomes translocate to the cell periphery upon overexpression of protrudin, and conversely, cluster perinuclearly upon protrudin depletion (Hong et al., 2017). Thus, we next tested whether recruitment of protrudin to lysosomes is TFEB-dependent. Again, we first induced nuclear translocation and thus activation of TFEB by rapamycin treatment of RT112 cells and visualized protrudin by immunofluorescence (**Fig. 3G**). Because protrudin is an ER-localized protein and only is found on lysosomes at ER-lysosome contact sites, we measured the fraction of protrudin that is found on LAMP1-positive lysosomes (**Fig. 3H**). We revealed that activation of TFEB significantly increased the fraction of protrudin found on lysosomes. Concomitantly, depletion of TFEB by siRNA in high-grade JMSU1 cells significantly reduced protrudin levels on LAMP1-positive lysosomes (**Fig. 3I, J**). Interestingly, protrudin gene expression was not up-regulated in high-grade cells (**Fig. S3D**), nor did the total protein level of protrudin change after rapamycin treatment in RT112 cells or when TFEB was targeted by siRNA in JMSU1 (**Fig. S3E, F**). This suggested that TFEB specifically regulated the recruitment of protrudin to lysosomes rather than its expression levels. Together our results indicate that lysosome positioning and protrudin recruitment on lysosomes in bladder cancer cells is under the control of TFEB.

### TFEB regulates phosphatidylinositol-3-phosphate levels on endomembranes in bladder cancer cells

Recruitment of protrudin to lysosomes is regulated by the binding of its FYVE domain to phosphatidylinositol-3-phosphate (PtdIns3P) found on endomembranes (Hong et al., 2017). We thus tested whether TFEB could regulate lysosomal PtdIns3P levels. We expressed the PtdIns3P-binding FYVE domain from the human homologue of the hepatocyte growth factor-regulated tyrosine kinase substrate Hrs, duplicated in tandem as an EGFP fusion construct (EGFP-FYVE) and monitored total level of this construct on LAMP1-positive lysosomes upon knock down of TFEB. PtdIns3P is found on early and late endosomes, thus as expected, EGFP-FYVE showed an endosomal staining but only partially colocalized with lysosomes (**Fig. 4A**). Silencing of TFEB by siRNA significantly decreased the fraction of EGFP-FYVE that was found on Lamp1-positve lysosomes in high-grade JMSU1 cells (**Fig. 4B**). Moreover, measuring the global cellular level of EGFP-FYVE showed a significant reduction on endomembranes after knock down of TFEB (**Fig. 4C, D**).

**Figure 4.**
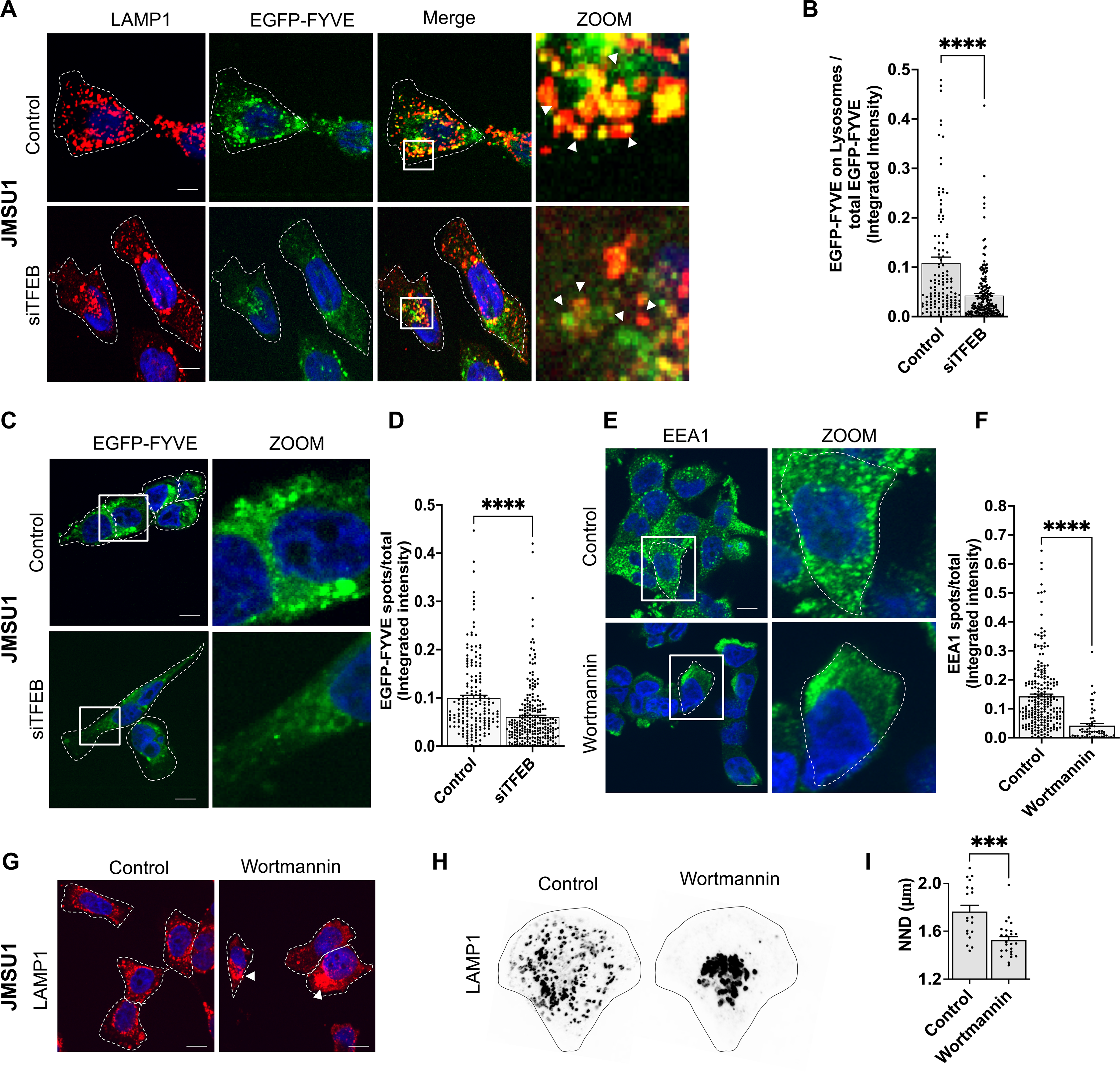
TFEB regulates phosphatidylinositol-3-phosphate levels on endomembranes in bladder cancer cells. **A**. Immunofluorescence staining of the lysosomal-associated membrane protein 1 (LAMP1/CD107a) (red) in control (siLUC) and siTFEB (72 h) treated JMSU1 cells transfected with EGFP-FYVE (green). Zoom shows the merged images of LAMP1 and EGFP-FYVE in white box. White arrow shows the colocalization between LAMP1 and EGFP-FYVE. Scale bars equal 10 μm. **B**. Quantification of EGFP-FYVE integrated intensity on lysosomes normalized to total cellular EGFP-FYVE, in 147 siLUC and 167 siTFEB treated JMSU1 cells; **** p<0.0001 in a Mann-Whitney U test, error bars are SEM. **C**. Representative images of control (siLUC) and siTFEB (72 h) treated JMSU1 cells expressing EGFP-FYVE. Zoom shows EGFP-FYVE in white box. Scale bars equal 15 μm. **D**. Quantification of EGFP-FYVE integrated intensity on segmented spots normalized to total cellular EGFP-FYVE, in 241 siLUC and 307 siTFEB JMSU1 cells; p<0.0001 in a Mann-Whitney U test, error bars are SEM. **E**. Immunofluorescence staining of early endosome antigen 1 (EEA1) in control (DMSO) and wortmannin (1 μM) treated JMSU1 cells. Zoom shows one single cell in white box. Scale bars equal 15 μm. **F**. Quantification of EEA1 integrated intensity on segmented spots normalized to total cellular EEA1, in 228 control and 56 wortmannin treated JMSU1 cells; **** p<0.0001 in a Mann-Whitney U test, error bars are SEM. **G**. Immunofluorescence staining of LAMP1 in control (DMSO) and wortmannin (1 μM) treated JMSU1 cells. White arrows show the perinuclear clustering of lysosomes. **H**. Representative images of lysosomes visualized by immunofluorescence staining against LAMP1 in micropatterned control and wortmannin (1 μM) treated JMSU1 cells. **I**. Nearest neighbor distance (NND in μm) between lysosomes in micropatterned in control (n=19) and wortmannin (n=25) JMSU1 cells; *** p < 0.0005 p-value in a Mann-Whitney U test, error bars are SEM.

To further validate the TFEB regulated recruitment of PtdIns3P-binding proteins to endosomal membranes, we analyzed EEA1, a well-studied FYVE containing protein. Consistent with protrudin, we found a significant increase of EEA1 on endomembranes upon treatment of low-grade RT112 cells with rapamycin and activation of TFEB (**Fig. S4A, B**). Addition of the protein translation inhibitor cycloheximide along with rapamycin treatment abolished the increase of EEA1 on endomembranes. Surprisingly, total EEA1 protein levels did not change under tested conditions (**Fig. S4A-C**) although EEA1 expression has been previously described to be under the control of TFEB (Nnah et al., 2019). No increase of endosomal EEA1 was observed in the first 4 h after rapamycin treatment (**Fig. S4D-F**). Conversely, gene silencing of TFEB in high-grade JMSU1 cells significantly decreased EEA1 levels on endosomes without affecting the total amount of EEA protein level (**Fig. S4G-I**).

Finally, we tested the role of endosomal PtdIns3P levels on protein recruitment and lysosome positioning. As expected, inhibition of phosphatidylinositol-3-phosphate kinases by wortmannin in high-grade JMSU1 cells strongly depleted EEA1 from endosomes (**Fig. 4E, F**) mimicking the phenotype of TFEB knock down (**Fig. S4G-I**). Moreover, wortmannin treatment induced the central clustering of lysosomes in JMSU1 cells, leading to a significant reduction of the NND of lysosomes (**Fig. 3E, F**). This showed that dispersion of lysosomes towards cell periphery requires endosomal PtdIns3P. Altogether, our results indicate that endosomal PtdIns3P levels dictate lysosomal positioning and are regulated by TFEB in high-grade bladder cancer.

## Discussion

Our study identifies and characterizes a novel cellular phenotype of aggressive malignancy in a cellular model of bladder cancer. We show that the lysosomal compartment is scattered to the cell periphery in all analyzed high-grade bladder cancer cells, a phenotype that we did not see in normal urothelial cells. This is different to the previously described expansion of the lysosomal compartment, characterized by an increase in volume or numbers of lysosomes in pancreatic ductal adenocarcinoma (PDA) and indicative of increased lysosome biogenesis (Perera et al., 2015). Moreover, lysosome positioning changes are correlated with changes in mTORC1 signaling that assembles on lysosomes. In high-grade cells, the classic mTORC1 substrate p70-S6K1 was less phosphorylated, and TFEB translocated to the nucleus, potentially due to either reduced phosphorylation by mTORC1 or increased dephosphorylation by the calcium-dependent phosphatase, calcineurin (Medina et al., 2015). Deregulation of mTORC1 signaling and TFEB hyper-activation parallels previous studies in other cancer types (Bar-Peled et al., 2013; Di Malta et al., 2017; Perera and Zoncu, 2016; Perera et al., 2019; Zoncu et al., 2011) and aligns with a genetic study of NMIBC that identified alterations in mTORC1 signaling in several bladder cancer subtypes (Hurst et al., 2017).

Peripheral dispersion of lysosomes has been previously reported in prostate cancer cells due to the acidification of the extracellular milieu (Steffan et al., 2009). Such a mechanism is unlikely in the case of bladder cancer cells, because all cells used in this study were grown in the same pH-buffering medium. Reports that TFEB regulated lysosome positioning (Medina et al., 2011; Willett et al., 2017), and the fact that peripheral lysosomes correlated with a hyperactivated TFEB phenotype (nuclear localization of TFEB-GFP), let us to investigate if TFEB regulated lysosome positioning in bladder cancer cells. We found that induction of nuclear TFEB by rapamycin in low-grade RT112 cells induced lysosomal dispersion. Conversely, knockdown of TFEB in high-grade JMSU1 cells with nuclear TFEB-GFP induced perinuclear clustering of lysosomes. Thus, our results confirm that lysosome positioning is under the regulation of TFEB in the bladder cancer model.

TFEB is a key transcription factor that orchestrates the expression of many genes involved in metabolism (Sardiello et al., 2009; Settembre et al., 2013) but also in intracellular trafficking of organelles (Nnah et al., 2019). Investigating the molecular mechanisms by which TFEB controls lysosome positioning, we discovered that TFEB regulates endosomal PtdIns3P levels that leads to enhanced recruitment of FYVE-containing proteins such as protrudin. Protrudin is known to bind to PtdIns3P on endosomes at membrane contact sites with the endoplasmic reticulum (ER) and to recruit the kinesin-1 adaptors FYCO1 to promote the microtubule-dependent translocation of endosomes to the cell periphery (Pedersen et al., 2020; Raiborg et al., 2015). However, several alternative pathways for anterograde lysosome trafficking have been described that all require endosomal PtdIns3P and could additionally be harnessed by cancer cells. The alternative kinesin-1 adaptor SKIP (also known as PLEKHM2) also contains three lipid-binding pleckstrin homology (PH) that conceivably could bind to lysosomal PtdIns3P. Moreover, KIF16B, a highly processive kinesin-3 family member that participates in the trafficking of endosomes along microtubules contains a PX (Phox homology) motif binding PtdIns3P (Pyrpassopoulos et al., 2017). Importantly, whereas we only observed a moderate change in the protein levels of the direct TFEB target EEA1 after knock down of TFEB, our data indicates that the upregulation of endosomal PtdIns3P levels is transcriptionally regulated. Indeed, increase of endosomal FYVE-protein recruitment was not obvious in the first 4 hours after nuclear TFEB induction and was abolished upon the protein translation inhibitor cycloheximide (**Fig. S4 A-F**). PtdIns3P formation depends on either the class II phosphatidylinositol 3-kinase (PI3K) PI3KC2A, or the class III PI3K Vps34 (or PIK3C3) (Burke, 2018). Although no experimental evidence currently shows that TFEB regulates the expression of class III PI3 kinase, it has been shown in skeletal muscles that TFEB overexpression induced the expression of several PI kinases subunits, for instance, PIK3CD, PIK3C2A (Mansueto et al., 2017). Moreover, TFEB is known to regulate genes involved in lipid catabolism in liver and skeletal muscle (Settembre et al., 2013), some via co-induction of peroxisome proliferator-activated receptor-γ coactivator1α (PGC1α) and peroxisome proliferator-activated receptors (PPAR). Because PIP metabolism is complex and dynamic, additional studies are required to reveal the specific mechanisms of PtdIns3P increase.

Interestingly, it has been shown that endosomal PtdIns3P levels regulate mTORC1 recruitment and signaling via amino acids and stimulation of class III PI3K/Vps34-mediated PtdIns3P synthesis (Gulati and Thomas, 2007). PtdIns3P also facilitates lysosomal recruitment of phospholipase D1 (PLD1) via its PX domain that produces phosphatidic acid, which triggers dissociation of the inhibitory DEPTOR subunit from mTORC1 (Song and Yoon, 2016). Additionally, the PtdIns3P3-phosphatase MTMR3 interacts with mTORC1, and overexpression of this enzyme inhibits mTORC1 activity (Hao et al., 2016). Finally, the formation of phosphatidylinositol 3,5-bisphosphate (PtdIns3,5P_2_) from PtdIns3P regulates mTORC1 via raptor (Bridges et al., 2012). As increased endosomal PtdIns3P levels globally activate mTORC1 that deactivates TFEB, we speculate that PtdIns3P could be part of a feedback loop in the mTORC1/TFEB signaling axis. Indeed, TFEB has been shown to feedback on mTORC1 (Nnah et al., 2019). Our current understanding is that nutrient status, pH and growth factors assemble a sophisticated machinery on the surface of lysosomes to integrate the different inputs upstream of mTORC1 (Ballabio and Bonifacino, 2020; Lawrence and Zoncu, 2019; Shin and Zoncu, 2020). Because PtdIns3P and several motor proteins/adapters are part of this machinery, mTORC1 signaling is coupled with lysosome positioning (Korolchuk et al., 2011). Our data are consistent with the following model: In high-grade bladder cancer cells, TFEB localization is mostly nuclear. Nuclear presence of TFEB and its transcription activity leads to an increase in PtdIns3P levels on different endomembranes, including lysosomes. This increase leads to the recruitment of FYVE-domain containing proteins such as EEA1 and protrudin and supports anterograde movement of lysosomes. The anterograde movement gives rise to the typical signature of peripheral lysosomes that we find in all studied high-grade bladder cancer cells. Peripheral lysosomes have been shown to recruit more mTORC1 and increase phosphorylation of downstream substrates (Hong et al., 2017; Korolchuk et al., 2011; Perera and Zoncu, 2016). This would allow a feedback control of TFEB by mTORC1. However, this seems not to occur in high-grade bladder cancer cells, because mTOR levels on lysosomes do not increase, and another mTORC1 substrate (S6K) shows less phosphorylation. Instead, the efficient calcium-dependent dephosphorylation of TFEB hinders its cytoplasmic translocation and control by mTORC1 in bladder cancer cells. Together, our results provide a mechanistic explanation to the characteristic cellular phenotype of lysosome dispersion in high-grade bladder cancer cells. Yet, further studies will be required to reveal in detail the deregulation of the mTORC1/TFEB axis in different bladder cancer cells. In addition to signaling, lysosome positioning has been implicated in the regulation of protease secretion/proteolysis (Pedersen et al., 2020), migration (Dozynkiewicz et al., 2012; Pu et al., 2015, 2016; Schiefermeier et al., 2014) and remodeling of the tumor environment through the release of exosomes (Hyenne et al., 2017). Thus, it is tempting to speculate that altered lysosome signaling could link dysfunctional cancer cell metabolism with cancer invasiveness.

In addition to revealing a novel cellular phenotype characteristic of cancer cells together with the underlying molecular mechanism, our results uncover a novel role of TFEB in regulating PtdIns3Ps levels on endosomes. Several studies have illustrated the crucial role of TFEB in regulating fundamental but distinct cellular processes such as endocytosis, lysosomal biogenesis and autophagy. Because these different compartments of the endolysosomal system retain their identities based on the lipid composition of their membranes and are regulated by PtdIns3P levels, our results conceptually clarify the role of TFEB as regulator of endosomal maturation.

## Material and methods

### Cell culture and treatments

Bladder cancer cells lines MGHU3, RT112, KU19-19, JMSU1, T24 and TCCSup were grown in RPMI-1640 medium (Life Technologies, Carlsbad, CA, USA), supplemented with 10% Fetal Bovine Serum (FBS; Eurobio, Courtaboeuf, France). Normal human urothelium (NHU) cells were from Jennifer Southgate (University of York, UK). NHU were grown in KSFMC medium according to (Southgate et al., 1994). For experiments with inhibitors, as per the experiment either the day after cell seeding or after transfection respective drugs were added for incubation time of 24 h or as indicated and cells were incubated, at 37°C. The concentration of inhibitors used were as follows: rapamycin (10 μM), wortmannin (1 μM, 2 h), ML-SI1 (20 μM, 3 h), BAPTA AM (10μM, 3 h) and cycloheximide (20 μg/mL). For starvation experiments, the day after cell seeding, the medium was removed and cells were washed once with EBSS (Earle’s Balanced Salt Solution) and incubated in EBSS for 4 or 24 h, as per the experiment, before lysate preparation or cell fixation with 4% PFA.

### Cell transfection

For RNA interference studies, 200,000 cells were transfected in 12 well plate with 25 pmol siRNA (siTFEB : ON-TARGETplus Human TFEB, L-009798-00-0005, Dharmacon™) using Lipofectamine RNAiMAX Transfection Reagent (1:200; Life Technologies). Cells were incubated 72 h at 37°C prior to further manipulation or drug treatment. Efficiency of gene silencing was verified by western blot of cell lysate after three days of transfection.

For plasmid transfection, 200,000 cells were transfected in a 12 well plate. Transfection was performed using Lipofectamine LTX with Plus reagent (Invitrogen) using 1 μg of plasmid. pEGFP-N1-TFEB plasmid was a gift from Shawn Ferguson (Addgene plasmid # 38119; http://n2t.net/addgene:38119; RRID:Addgene_38119n (Roczniak-Ferguson et al., 2012)) or EGFP-2X FYVE plasmid (kind gift from B. Payrastre, Toulouse). 48 h post transfection, cells were trypsinized and transferred to sterilized coverslips (12 mm) in 1 mL medium in 12 well plate. Cells were fixed with 4%PFA 72 h after transfection and used for immunofluorescence and imaging.

### Micro-array analysis

Micro array data were analyzed with R (3.5.2). The annotation was performed using affy package (1.58.0) with a custom CDF (Chip Description File) from brain array (huex10st, genome version 23). Normalization was done with RMA algorithm using affy library (Gautier et al., 2004) and batch effect corrected with ComBat (Johnson et al., 2007). The PCA was computed from these normalized and corrected data.

### Micropatterned coverslips preparation and cell seeding

Micropattern production was as previously described (Duong et al., 2012; Schauer et al., 2010a) using photo-lithography methods. Briefly, coverslips were coated with Poly-L-Lysine(20)-grafted[3.5]-Polyethyleneglycol(2) (PLL-g-PEG) from SuSoS (Dübendorf, Switzerland) at a final concentration of 0.1 mg/mL in 10 mM HEPES (pH 7,3) solution. Coverslips were exposed to deep UV during 5 min using a photomask containing arrays of crossbows (37 μm diameter, 7 μm thick). Prior to cell seeding, the patterned surface was incubated for 1 h with a mixture of 50 μg/mL fibronectin (Sigma-Aldrich, St. Louis, MO, USA), 5 μg/mL concanavalin A (Sigma-Aldrich, St. Louis, MO, USA) and 1 μg/mL fibrinogen–Cy5 (Invitrogen). Cells were seeded on micropatterns in RPMI medium supplemented with 20 mM HEPES (Life Technologies) for 4 h prior the experiment.

### Invasion assay

Cells were trypsinized and 10^4^ cells/ml were re-suspended in RPMI medium containing 10% FBS and 1% Penicillin-Streptomycin (Life Technologies). Then 100 μl of cell suspension was plated in 48-well plates coated with 1% agarose (Life Technologies) and incubated for 3 days. In each well, a spheroid was formed from 10^3^ cells. Next, the spheroids were plated on Lab-Tek chambers (Sigma), in a mixture of collagen I from rat tail (Corning) at a final concentration of 2 mg/ml, PBS, sodium hydroxide (NaOH) and serum-free medium. The spheroids were monitored for 5 consecutive days by using an inverted Leica microscope (Wetzlar, Alemanha) equipped with camera device using 4x objective.

### Immunofluorescence

Cells were fixed with 4 % formaldehyde for 15 min at room temperature, washed three times with PBS and permeabilized in PBS containing 0.2% BSA and 0.05% saponin. Cells were then incubated with the primary antibodies (mouse monoclonal antibody against Lamp1/CD107a (555798, BP Pharmingen™), rabbit mAb against mTOR (7C10, #2983, Cell Signaling Technology), EEA1 (610456, BD Biosciences), protrudin / ZFYVE27 (12680-1-AP, Proteintech) and Alexa Fluor 488, or Alexa Fluor 647 or Cy3-coupled secondary antibodies (Jackson ImmunoResearch) for 1h. Actin was visualized by FluoProbes 547H (557/572nm) coupled Phalloïdin (Interchim) and nuclei with 0.2 μg/mL 4’,6-diamidino-2-phenylindole (DAPI; Sigma-Aldrich). Coverslips were mounted in Mowiol (Sigma-Aldrich).

### Western Blot

250,000 cells were seeded in a 12 well plate one day prior to the experiment. Drug treatments or knock-down experiments were performed as mentioned before. Equal volumes of lysate from each cell line was loaded on a 10% or 12% polyacrylamide gel, resolved by SDS-PAGE and electrotransferred to nitrocellulose membranes. Membranes were incubated with primary antibodies at 4°C overnight: Phospho P-70 (Thr389)-S6K (CST: 9205S, 1:1000 in 5% BSA in TBST), P-70 S6K (CST: 9202S, 1:1000 in 5% milk in TBST), GAPDH (Sigma: G9545, 1:10,000 in 5% milk in TBST), EEA1(610456, BD Biosciences, 1:500 in 5% milk in TBST), protrudin (ZFYVE27, Proteintech 12680-1-AP) and species specific HRP secondary antibodies (1:10,000) for 1 hour at room temperature, following ECL western blotting substrate.

### Image acquisition

Images for immunolabelled cells on micropatterns were acquired with an inverted wide field Deltavision Core Microscope (Applied Precision) equipped with highly sensitive cooled interlined charge-coupled device (CCD) camera (CoolSnap Hq2, Photometrics). Z-dimension series were acquired every 0.5 μm.

Images for non-pattered immuolabelled cells were acquired with a spinning disk confocal microscope (Inverted Eclipse Ti-E (Nikon) + spinning disk CSU-X1 (Yokogawa) integrated with Metamorph software by Gataca Systems). Cells were imaged as Z-stacks with 0.2 μm distance and 12 μm total height.

### Image processing and analysis

For cells on micropatterns, several tens of single cell images were aligned using the coordinates of the micropattern (determined on ImageJ (Bethesda, MD, USA) as previously described (Grossier et al., 2014; Schauer et al., 2010b). To extract the 3D spatial coordinates of lysosomes, images were segmented with the multidimensional image analysis (MIA) interface on MetaMorph (Molecular Devices, Sunnyvale, CA, USA) based on wavelet decomposition. The coordinates of the segmented structures were processed for density estimation programmed in the ks library in R according to (Schauer et al., 2010b). For visualizing kernel density estimates, probability contours were visualized using the extension libraries mvtnorm, rgl, and miscd.

Levels of lysosome dispersion in non patterned MGHU3, RT112, KU19-19 and JMSU1 cells were measured using statistical inertia (=averaged squared distance to the center of mass). To control for variations in cell size differences, normalization to cell size has been applied. Lysosome coordinates have been divided by the coordinates of the center of the mass (setting the center mass at x=1, y=1). This quantifies the dispersing of the lysosome structures independently of homogeneous dilations due to cell size. To test statistical significance, a Kruskal-Wallis test with Dunn post-hoc test with Sidak correction for multiple comparisons has been applied.

Image analysis for the figures (2B, 3H, J, 4 B,D,F and S4B, E, H) was done using CellProfiler (version: 3.1.9) on one Z-plane of the images. The pipelines for different analysis were prepared as follows:

*To detect the total and membrane bound intensities of protein of interest (labelled as total integrated intensity or spots/total, respectively, in the figures) or intensities of co-localized proteins the pipeline was created as follows:*

Step 1: Module ‘*EnhanceorSuppressFeatures*’ was applied to channels where the objects needs to be segmented, either to obtain their intensities or objects for the intensities of co-localized proteins, to get sharp and defined objects which makes segmentation easier (for eg. On channels with LAMP1 or EEA1 or GFP-FYVE).

Step 2: Nucleus was identified in the DAPI channel using the ‘*IndentifyPrimaryObject*’ module

Step 3: Module ‘*IndentifyPrimaryObject*’ was used again on the images obtained from Step 1 to segments objects whose measurements are required (such LAMP1, EEA1, EGFP-FYVE)

Step 4: Cells were segmented using the ‘*IdentifySecondaryObject*’ module with nucleus as the ‘primary object’ (identified in step 2) and using phalloidin or another cytoplasmic protein channel to recognize the cell boundaries.

Step 5: Module ‘*RelateObjects’* was used to relate the objects obtained in step 3 to each cell obtained in Step 4. Output of this channel was saved as another object which gives the objects of protein of interest per cell.

Step 6: Objects from step 3 were masked on the channel whose co-location or membrane bound fraction had to be calculated using the ‘*MaskImage*’ module. (for eg: to calculate EGFP-FYVE on lysosomes in Fig. 4B, Lysosomes were segmented in step 3 and the output objects were masked on EGFP-FYVE channel or to calculate membrane bound EGFP-FYVE, segmentation of EGFP-FYVE objects from step 3 was masked on EGFP-FYVE channel)). Output of this step was saved as a new image in the pipeline.

Step 7: ‘*MeasureObjectIntensity*’ module was used to obtain total ‘per cell intensity’ and ‘intensity on spots’ of protein of interest. Intensities were picked from images from step 6 and raw images of channel of interest using cells from step 4 as the objects.

Step 8: Cell size was obtained using the module ‘*MeasureObjectSizeandShape*’ on the cells segments in Step 4 as the objects

Step 9: Finally, all the measurements were exported to the excel sheet using the module ‘*ExporttoSpreadsheet*’

Step 10: The final values were exported to a csv file named ‘cell’. This file had the values of cell size (in pixels), total intensity of protein of interest per cell, intensity of protein of interest on spots and intensity of co-localized protein on the object of interest (eg: GFP-FYVE on lysosomes). Integrated intensities were used for the analysis and to plot the graphs.

### Statistical analysis

The statistical analysis of endolysosome volume, number and normalized NND was performed with R (3.6.0). For NND analysis, the centroids distance between structures was calculated from a constant number of lysosomes that was randomly sampled from each cell. Therefore, variation in NNDs cannot be imputed to variation in the number of lysosomes but to bona-fide variation of their spatial organization. The statistical analysis was a Kruskal-Wallis test with Dunn test for multiple comparisons correction.

For all experiment, a large number of cells were monitored from 3 to 6 independent experiments. Two-sided Mann-Whitney U test were performed for 2 conditions comparisons. For multiple comparisons, a Kruskal-Wallis has been used with Dunn’s test for multiple comparisons. Additionally, to compare the global distribution of cell population, χ² tests were performed (R function “chi-square()”) and Benjamini-Hochberg multiple comparison correction has been applied. For the statistical analysis on the data from CellProfiler, Prism was used. Mann-Whitney U test was applied for the two conditions comparison or Kruskall-Wallis test with Dunn test for multiple comparison.

## Supporting information

SuppFigures1-4

## Acknowledgements

We thank Jennifer Southgate (University of York, UK) for her gift of NHU cell extracts and Clémentine Krucker and Yann Neuzillet (Foch Hospital, France) for deriving these cells. We thank Elodie Chapeaublanc for help on the bioinformatics and Oliver Kepp for critical reading of the manuscript. The authors greatly acknowledge the Cell and Tissue Imaging Facility (PICT-IBiSA @Burg and @Pasteur) and the Nikon Imaging Center at Institut Curie (Paris) that are member of the French National Research Infrastructure France-BioImaging (ANR10-INBS-04). This project was supported by grants from the European Union’s Horizon 2020 research and innovation program under the Marie Skłodowska-Curie grant agreement n° 666003 to PM; Capes/ Ciência sem Fronteiras/ Process (9121137) to CDBS; Grants from Fondation ARC pour la recherche sur le cancer to KS and HL, Institut Curie SIRIC grant to KS, Agence Nationale de la Recherche (#2010 BLAN 122902), the ITMO Nanotumor grant to KS, the Centre National de la Recherche Scientifique and Institut Curie. The Goud team is member of the Labex Cell(n)Scale (11-LBX-0038) and the Idex Paris Sciences et Lettres (ANR-10-IDEX-0001-02 PSL). The Molecular Oncology team (FR) is supported by La Ligue Contre le Cancer (Equipe labellisée program).

## Supplemental figure legends

**Figure S1. High-grade cancer cell lines are specifically characterized by scattered, peripheral positioning of lysosomes**

**A**. Average intensity projections of the actin cytoskeleton visualized by phalloidin of n cells of normal human urothelium (NHU) and bladder cancer cell lines RT4 (ATCC^®^ HTB-2™), MGHU3 (Lin et al., 1985), RT112 (Marshall et al., 1977), KU19-19 (Tachibana et al., 1995), T24, TCCSup (Nayak et al., 1977), JMSU1 (Morita et al., 1995). Scale bar equals 10 μm. **B**. Correlation analysis between average endolysosomal volume and average numbers per cell shows a weak (R²=0.19) but significant association; p-value < 0.001 in a t-test for correlation. **C**. Principal component analysis of transcriptome data of normal human urothelium (NHU) cells and the bladder cancer cell lines RT4, MGHU3, RT112, KU19-19, T24, TCCSup and JMSU1. **D**. Average day of invasion from spheroids into collagen matrix of MGHU3 (n=13), RT112 (n=9), KU19-19 (n=5), and JMSU1 (n=8), and representative images of 3D spheroids from KU19-19 (upper panel) and JMSU1 (lower panel) at 1 day after matrix embedding. White arrow indicates invasion of collagen matrix by escaping cells. Scale bar equals 500 μm. **E**. Schematic representation of the analysis of endolysosome distribution in classical cell culture conditions (see **F**). **F**. Normalized lysosome dispersion in non patterned MGHU3, RT112, KU19-19 and JMSU1 cells based on statistical inertia measurement (=averaged squared distance to the center of mass normalized to cell size) for n>60 cells per cell line analyzed, **** p < 0.0001 in a Kruskal-Wallis test with Dunn‘s test for multiple comparison.

**Figure S2. Altered lysosomes correlate with changes in the mTORC1-TFEB nutrient signaling pathway**

**A**. Quantification of total p70-S6 Kinase 1 levels from n=3 Western Blot experiments in MGHU3, RT112, KU19-19 and JMSU1 (see also Figure 2C). Error bars show SEM. **B**. Western Blot analysis of phosphorylated p70-S6 Kinase 1 (P-p70-S6K1 Thr389) and GAPDH loading control in MGHU3, RT112, KU19-19 and JMSU1 cells in control conditions (full media) and after treatment with rapamycin at 10 μM for 2 h or grown under starvation in EBSS (Earle’s Balanced Saline Solution) for 4 h. **C**. Normalized Log2 RNA expression levels of housekeeping gene (GAPDH) and TFEB regulated genes (RRAGD and TSC1) in NHU, MGHU3, RT112, KU19-19 and JMSU1. **D**. Representative images of JMSU1 cells transfected with TFEB-EGFP for 72 h and treated with NPC1 inhibitor U18666A (10 μM, 2 h, 37°C). Scale bars are 10 μm. **E**. Quantification of the nuclear fraction of the total mean TFEB-EGFP fluorescent intensity in in control and U18666A treated JMSU1 cells (for n>15 cells in each condition). ns is p >0.1; Mann-Whitney test. Data are depicted as mean ± SD.

**Figure S3. Lysosome positioning changes are under the control of TFEB in bladder cancer cells**

**A**. Western blot analysis of siTFEB (72 h, with siRNA pool) in JMSU1 cells and quantification of TFEB levels normalized to GAPDH. Error bars are SEM of 7 independent experiments. **B**. Western blot of TFEB knockdown with individual siTFEB RNAs (72 h). **C**. Immunofluorescence staining against the lysosomal-associated membrane protein 1 (LAMP1, CD107a) in JMSU1 cells after TFEB knockdown with individual siTFEB RNAs (72 h). White arrows show the perinuclear clustering of lysosomes. Scale bar is 15 μm. **D**. Normalized Log2 RNA expression levels of protrudin (ZFYVE27). **E**. Western blot analysis of protrudin in RT112 cells in control (DMSO) and rapamycin (10 μM, 24 h) treatment conditions. **F**. Western blot analysis of protrudin in JMSU1 cells in control (siLUC) and siTFEB (72 h) conditions.

**Figure S4. TFEB regulates phosphatidylinositol-3-phosphate levels on endomembranes in bladder cancer cells**

**A**. Representative images of EEA1 staining in RT112 cells in control (DMSO), rapamycin (10 μM, 24 h) and cycloheximide (20 μg/mL, 24 h) conditions. Zoom shows one single cell in white box. Scale bars are 15 μm. **B**. Quantification of EEA1 integrated intensity on segmented spots normalized to total cellular EEA1, in control, rapamycin and cycloheximide treatment conditions in 234 control, 245 rapamycin and 201 cycloheximide treated RT112 cells; ****p<0.0001, ***p<0.001 in a Kruskal-Wallis test with Dunn’s test for multiple comparison, error bars are SEM. **C**. Western blot analysis of EEA1 in RT112 cells in control, rapamycin and cycloheximide treatment conditions. **D**. Representative images of EEA1 staining in RT112 cells in control (DMSO) and rapamycin (10 μM, 4 h) conditions. Zoom shows one single cell in white box. Scale bars are 15 μm. **E**. Quantification of EEA1 integrated intensity on segmented spots normalized to total cellular EEA1, in control and rapamycin conditions in 332 control and 330 rapamycin treated RT112 cells; ns p>0.1 in a Mann-Whitney test, error bars are SEM. **F**. Western blot analysis of EEA1 in RT112 cells in control and rapamycin (4 h) conditions, error bars are SEM from 3 independent experiments. **G**. Representative images of EEA1 staining in JMSU1 cells in control (siLUC) and siTFEB (72 h) conditions. Zoom shows one single cell in white box. Scale bars are 15 μm. **H**. Quantification of EEA1 integrated intensity on segmented spots normalized to total cellular EEA1, in control and siTFEB conditions in 94 siLUC and 212 siTFEB JMSU1 cells; ** p<0.01 in Mann-Whitney test, error bars are SEM). **I**. Western blot analysis of EEA1 in JMSU1 cells in control and siTFEB (72 h) conditions, error bars are SEM from 9 independent experiments.

