## Supplementary figures and images for "Transcription Factor EB regulates phosphatidylinositol-3-phosphate levels on endomembranes and alters lysosome positioning in the bladder cancer model"

### SuppFigures1-4

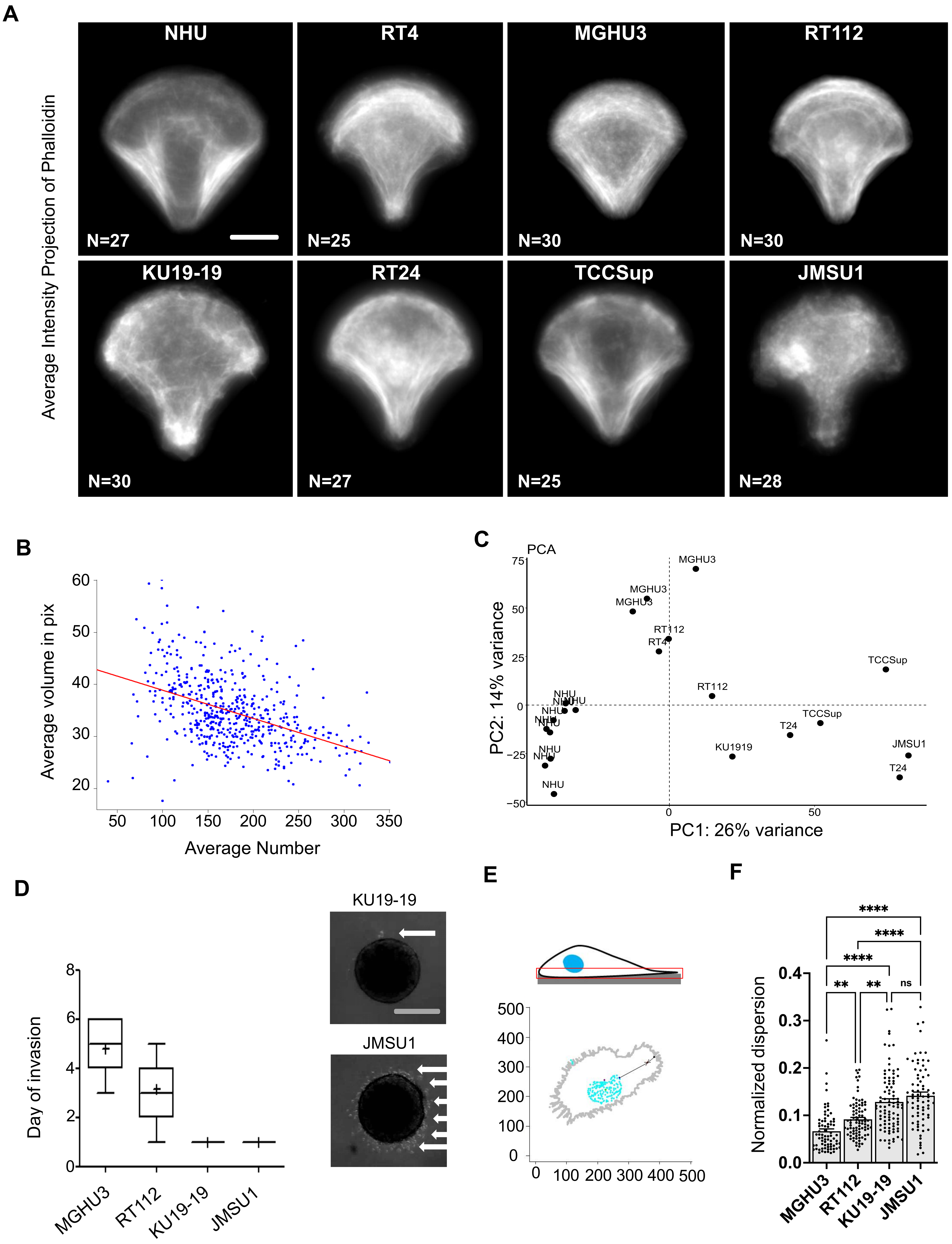


*Supplementary Figure 1*

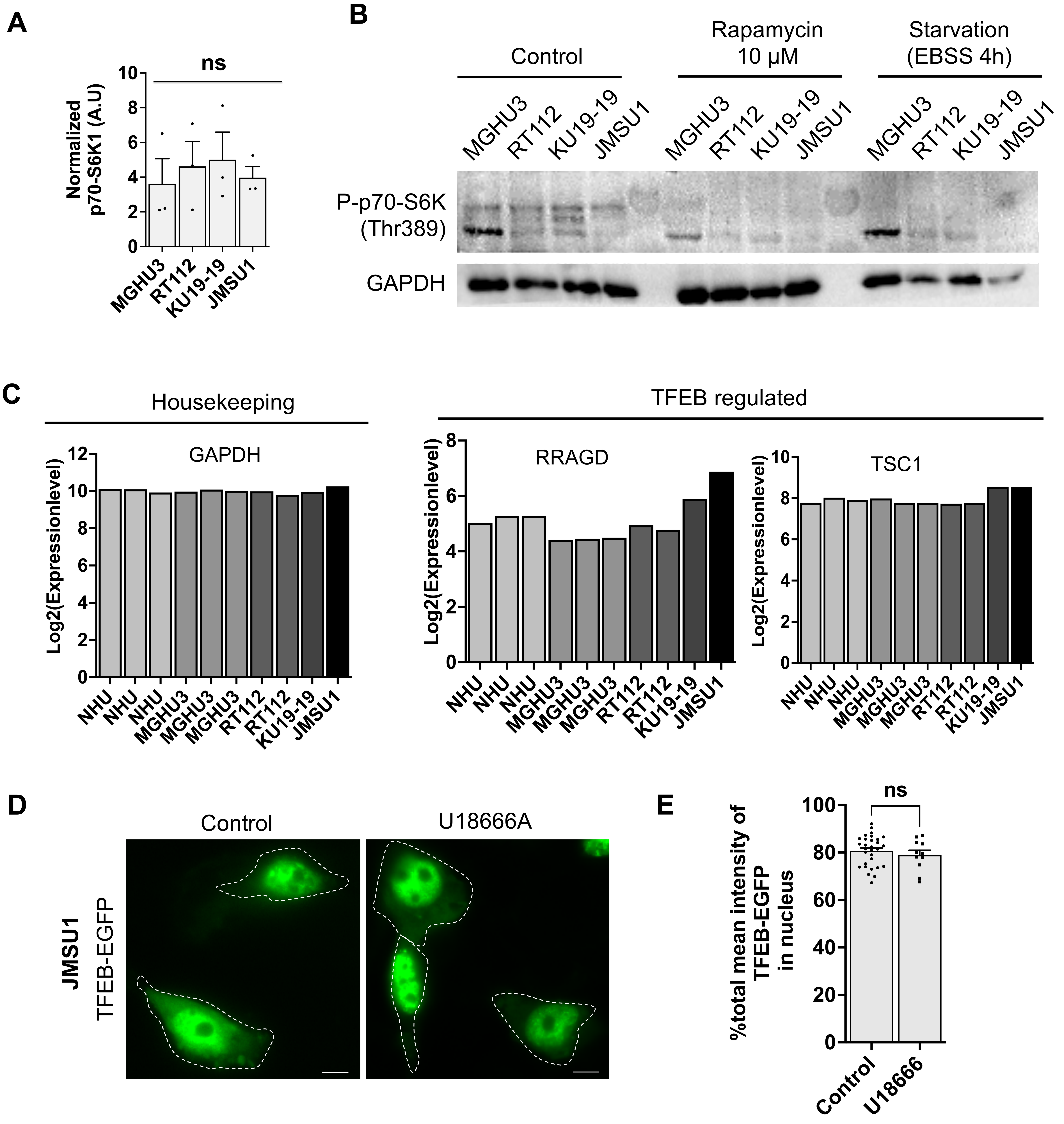


*Supplementary Figure 2*

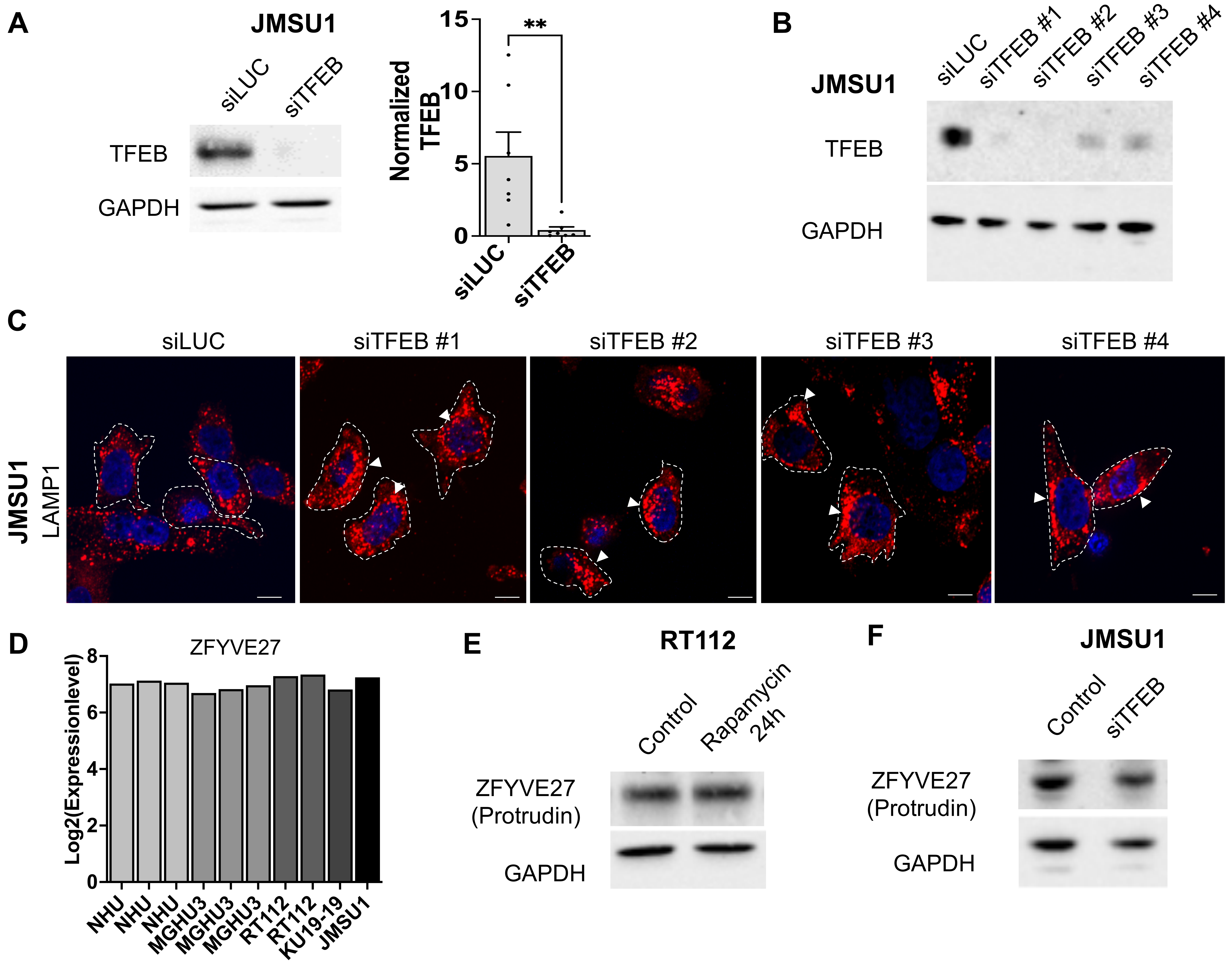


*Supplementary Figure 3*

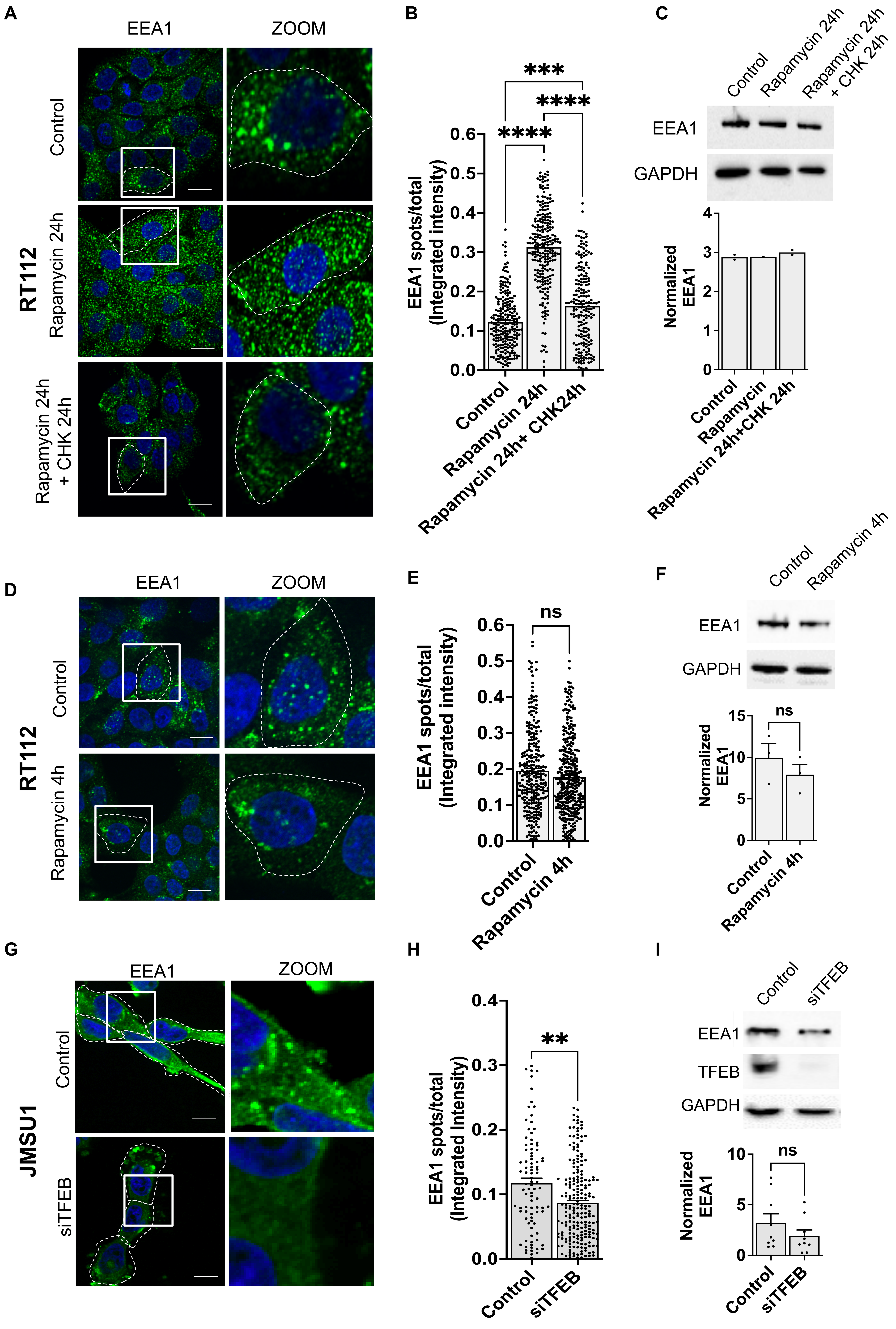


*Supplementary Figure 4*
